# Cycles of transcription and local translation support molecular long-term memory in the hippocampus

**DOI:** 10.1101/2021.10.29.466479

**Authors:** Sulagna Das, Pablo J. Lituma, Pablo E. Castillo, Robert H. Singer

**Author notes:** **Footnote:**.

## Abstract

Long-term memory requires transcription and translation of activity-regulated genes. Many of these are immediate early genes (IEGs) with short-lived mRNAs and proteins, decaying rapidly after stimulation. It remains unknown how an IEG with rapid mRNA and protein turnover can impact long-lasting changes at the synapses. Using fluorescently tagged endogenous *Arc*, an IEG important for memory consolidation, we performed high-resolution imaging of transcription and translation in individual neurons to identify the long-term gene dynamics after stimulation. Unexpectedly, once induced, *Arc* underwent transcriptional reactivation often at the same allele. Cycles of transcription were coordinated with localized translation. This cyclical regulation of an IEG, dependent on protein synthesis, reactivates subsequent transcription for maintaining mRNA supply to dendrites. The ensuing *Arc* mRNAs were preferentially localized at sites marked by previous Arc protein, thereby consolidating local “hubs” of dendritic Arc. These findings revealed the spatio-temporal dynamics of transcription-translation coupling of an IEG and provide a mechanism by which short-lived synaptic proteins can be sustained over the long-time scales of memory consolidation.

## Introduction

The ability to learn new information and store it for long periods ranging from hours to days and even a lifetime is one of the most remarkable features of the brain. Memory storage or consolidation requires modifications of synaptic connectivity and stabilization of these changes (Asok et al., 2019; Bailey et al., 2015). The molecular basis of long-term memory involves both transcription and translation (Alberini and Kandel, 2014; Wang et al., 2009), with growing evidence indicating the role of local translation (Richter and Klann, 2009; Sutton and Schuman, 2006). However, we still lack a detailed understanding of how transcription and translation are coordinated in space and in time to influence long term memory storage.

A class of genes termed as immediate early genes (IEGs) have established roles in transducing activity/experiences into long lasting molecular changes in neurons for memory (Plath et al., 2006; Tischmeyer and Grimm, 1999; Yap and Greenberg, 2018). Despite the importance of IEGs in memory storage, our understanding of how neuronal activity regulates IEG expression mostly focusses on the immediate early stages (within the first 2 hours after activity). Intriguingly, IEG mRNAs and proteins are characterized by short half-lives between 30-60 min (Rao et al., 2006; Tyssowski et al., 2018), that are inconsistent with the time frames of long-term memory.

Therefore, we attempted to resolve this conundrum by high-resolution imaging of transcription and translation dynamics of an IEG to determine its long-term regulation after initial activity. The Activity Regulated Cytoskeletal Associated (*Arc*) gene is an essential IEG for long-term memory and implicated in several neurodegenerative and cognitive disorders (Bramham et al., 2010; Guzowski et al., 2000; Plath et al., 2006). It is a unique multifunctional IEG possessing both synaptic and nuclear functions (Nikolaienko et al., 2018), and regulates different forms of synaptic plasticity: long term potentiation (LTP), long term depression (LTD), and homeostatic plasticity (Chowdhury et al., 2006; Messaoudi et al., 2007; Plath et al., 2006; Shepherd and Bear, 2011; Shepherd et al., 2006). A precise temporal regulation of Arc expression is critical for normal cognitive functions (Wall et al., 2018), and disrupted levels observed in various neurological disorders like Alzheimer’s (Wu et al., 2011), Angelman (Greer et al., 2010), FXS (Park et al., 2008), and schizophrenia (Fromer et al., 2014; Purcell et al., 2014). The prevailing view of *Arc* regulation is mostly derived from studying a short temporal window after neuronal stimulation and behavioral tasks (Guzowski et al., 1999; Steward et al., 1998; Tyssowski et al., 2018), although sustenance of Arc proteins in memory consolidation and recall have been shown (Messaoudi et al., 2007; Nakayama et al., 2015; Ramirez-Amaya et al., 2005). *Arc* mRNA is a natural target for non-sense mediated decay (Giorgi et al., 2007), and the protein on the other hand undergoes rapid degradation by ubiquitination (Soule et al., 2012). Given the rapid molecular turnover of Arc (mRNAs and protein half-lives approximately 60 min) (Das et al., 2018; Rao et al., 2006; Soule et al., 2012), how their levels are sustained to mediate long-lasting changes at the synapses is unknown. To address this question, we performed high resolution imaging of *Arc* mRNAs and proteins in hippocampal neurons over several hours after a brief stimulus. A knock-in Arc-PBS mouse, where the endogenous *Arc* gene was tagged with stem loops and detected by a fluorescent binding protein (Das et al., 2018), allowed us to follow *Arc* mRNAs from synthesis to transport and decay with high spatial and temporal resolution. The tagged *Arc* behaves similarly to the untagged WT gene in transcriptional output, mRNA half-lives and protein production, as well as in spatial memory tasks (Das et al., 2018).

We identified a unique feature of *Arc* gene regulation that helps to reconcile the inconsistencies between its mRNA/protein half-lives and its critical function in long term memory storage. Transcription was reactivated beyond the immediate early (IE) phase without additional neuronal stimulation, creating a supply chain of mRNAs, which are then available to undergo translation in the dendrites. Indeed, cycles of transcription were coordinated with distinct phases of local Arc protein synthesis. Importantly, translation occurred in “hotspots” along the dendrites, thereby forming local hubs of Arc protein. Over time, these hubs were maintained by subsequent cycles of mRNA synthesis, localization, and translation. By uncovering how activity driven IEG expression is maintained, and the crosstalk between transcription and translation, we provide a possible mechanism by which short-lived mRNAs and proteins with roles in synaptic plasticity could lead to long-lasting changes at the synapses required for memory consolidation.

## Results

### Arc gene undergoes transcriptional reactivation beyond the immediate early phase

While most studies focused on the immediate early (IE) phase of *Arc* transcription (within 60 min post stimulation), we investigated the long-term dynamics of the gene after an initial stimulus. Real-time imaging of transcription in hippocampal neurons from Arc-PBS mice was performed for four hours after stimulation triggered by tetrodotoxin withdrawal (TTX-w) (**Fig 1A**). The IE activation and shutdown was followed by a second transcriptionally active (“ON”) state, indicating a biphasic nature of *Arc* transcription. Since transcription was re-initiated in the same neuron after a prolonged OFF-period, we refer to this as “reactivation” (**Fig 1B, C**; **Movie S1**, **S2**). The average intensity trace of transcribing alleles from multiple neurons revealed that the two cycles of transcription were separated by an OFF period of 72 ± 9.3 min (**Fig 1D**). We classified the transcriptional dynamics beyond the IE-phase (100 mins post stimulation) into three categories-sustained, reactivated, and delayed (*de novo*) transcription. Transcription was considered “sustained” when any one of the alleles was active till 100 mins, “reactivated” when the transcription was re-initiated ≥30 mins after IE-phase, and “delayed” when *de novo* onset occurred after 90min. Notably, a significant 59.5 ± 3.1 % of transcription observed in the second cycle was a result of reactivation (**Fig 1E**), with an average onset time of 154.3 ± 4.2 min (avg ± SEM) (**Fig 1F**). A heat map of intensities from individual alleles in the neurons undergoing reactivation revealed that reactivation occurred often at the same allele in the population (**Fig S1A**). Comparison of two different stimulation conditions (TTX-w and chemical LTP) revealed that ∼60% of the transcriptionally active neurons from the IE-phase underwent reactivation in either case (**Fig S1B**).

**Fig. 1:**
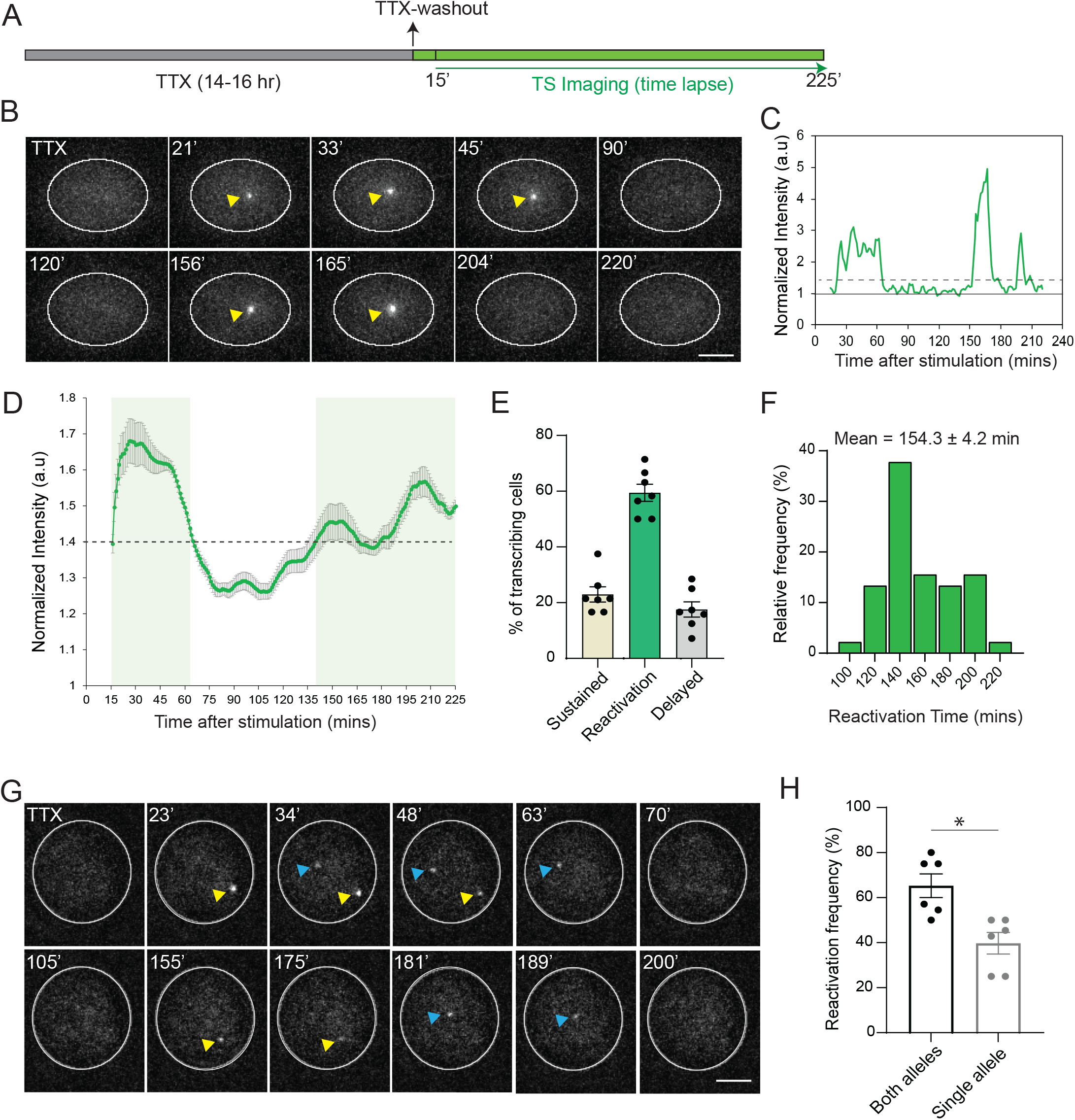
Long-term imaging of *Arc* transcription after stimulation. (**A**) Schematic of the stimulation paradigm, and the imaging time course. (**B**) Representative images showing the PCP-GFP labeled nucleus of a single neuron. Yellow arrows indicate transcribing Arc alleles. (**C**) Intensity trace of the transcribing allele in B. Solid line shows normalization to nuclear background. Dashed line indicates threshold for inclusion criteria as active transcription. (**D**) Average intensity trace from individual alleles of multiple neurons. Shaded areas indicate transcriptional activity above threshold (n= 58 neurons from 6 independent experiments). (**E**) Graph indicates the percentage of cells displaying different transcriptional states beyond the IE-phase (100 mins post stimulation). (n= 92 neurons, 7 independent experiments, each circle represents one experiment). (**F**) Frequency distribution of reactivation onset times (n=45 neurons, 6 independent experiments). (**G**) Representative images showing IE- and reactivation of both alleles albeit with different onset times. Arrows indicate transcribing alleles, yellow indicates first allele, and blue points to the second allele. (**H**) Frequency of reactivation from both alleles or from a single *Arc* allele (n=45 neurons, 6 independent experiments, each circle represents one experiment). Scale bar is 5 microns. Error bars indicate SEM.

The probability of allelic reactivation was determined, and a significantly higher frequency for both alleles was observed (62% for both vs 38% for single, p < 0.05, paired t-test) (**Fig 1G-H**, **Movie S2**). Reactivation was the result of an initial stimulation since stochastic transcriptional bursts from neurons undergoing basal network activity in the absence of TTX did not exhibit distinct cycles (**Fig S1C**). Moreover, transcription of *β-actin*, a constitutive housekeeping gene did not display obvious reactivation under TTX-w (**Fig S1D**), suggesting that the cycling is gene-specific with precise temporal regulation. Next, the output from the two transcriptional cycles were measured, and a significant reduction in nascent mRNA yield during reactivation compared to the IE stage was observed (**Fig S2A, B**). A similar decrease in the ON-duration during the reactivation was observed (**Fig S2C**), supporting that *Arc* transcriptional output was modulated primarily by the burst duration, which decrease over time leading to a possible dampening of the cycles. Thus, long-term imaging in individual neurons revealed the dynamics of *Arc* transcription beyond the IE phase and identified reactivation in a subset of neurons without any further stimulation.

### Optogenetic stimulation triggers transcriptional reactivation in cultures and in acute hippocampal slices

The long-term transcriptional response of individual neurons to a specific stimulus strength was performed using optical stimulation of single cells by expressing channel rhodopsin ChR2 in the neurons from the Arc-PBS mouse and monitoring of transcription over time. Selective stimulation of the soma was performed (**Fig 2A**) with trains of 20 Hz pulses (25 pulses/train) separated by two minutes (**Fig 2B**). Measurements of nuclear Ca^2+^ used as a read-out for activity displayed robust increases post-stimulation (**Fig 2C**). After opto-stimulation, biphasic transcription was observed in individual neurons (**Fig 2D**). The reactivation probability was 54.2 ± 4.2 %, with average onset time of 146.4 ± 7.1 min (**Fig 2E-F**), comparable to that observed in global stimulation (TTX-w, cLTP). A similar decrease in the transcriptional output due to lower ON-duration in the reactivation phase compared to the IE phase was also observed (**Fig S2D, E**).

**Fig 2:**
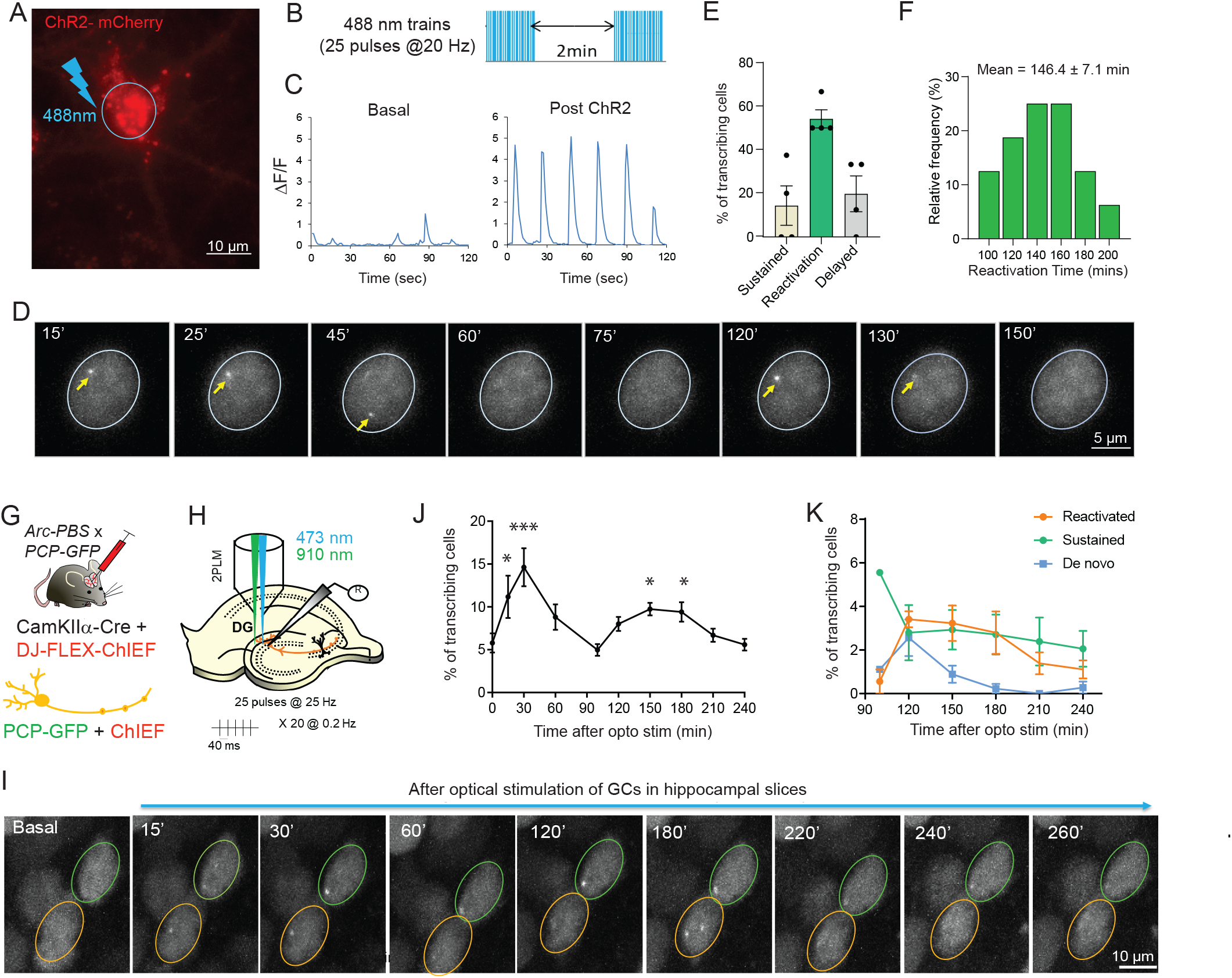
Reactivation of *Arc* transcription after optical stimulation in cultures and in tissue. (**A**) Hippocampal neurons from Arc-PBS animals were infected with lentiviruses expressing ChR2-mCherry. The soma of mCherry-positive neurons were stimulated with trains of 488 nm light using a pinhole (blue circle). (**B**) Stimulation paradigm for triggering activity. (**C**) Nuclear Ca^2+^ measurements in basal and after optical stimulation. (**D**) Representative images showing the PCP-GFP labeled nucleus of a single neuron after optical stimulation. Yellow arrows show transcribing Arc alleles. Scale bar 10 μm. (**E**) Percentage of cells displaying different transcriptional states after IE-phase (100 min after optical stimulation) (n = 27 neurons from 4 independent experiments). (**F**) Frequency distribution of reactivation onset times after optical stimulation of ChR2-expressing neurons. (**G**) Approach for Cre-dependent expression of PCP-GFP and ChIEF in the granule cells by injecting a cocktail of two viruses-CaMKIIα-Cre and DJ-FLEX-ChIEF to the dentate gyrus of Arc-PBS x PCP-GFP mice. (**H**) Stimulation paradigm for ChIEF (473nm) and two-photon imaging of GCs (910 nm illumination). (**I**) Representative images of two GC nuclei displaying transcription after optical stimulation. Orange outline shows neuron with transcriptional reactivation, green outline displays sustained activation. Scale bar 10 μm. (**J**) Total percentage of transcribing GC neurons after optical stimulation revealed two cycles (14.6 ± 2.2 % at 30 min, 9.7 ± 0.74 % at 150 min, 9.1 ± 0.9 % at 180 min, *** p = 0.003 at 30 min, *p = 0.01 at 150 min, *p= 0.04 at 180 min, compared to baseline 0 min, one-way ANOVA). (**K**) Distribution of the different transcriptional states in GC nuclei in the second cycle (100 min post stimulation). n = 5 slices, 5 animals (J, K). Error bars indicate SEM. *** denotes p < 0.005, * denotes p < 0.05.

To visualize the long-term transcriptional behavior of the *Arc* gene in tissue, an optogenetic stimulation approach was developed and transcription was monitored in real time in acute hippocampal slices from Arc PBS animals crossed with PCP-GFP transgenic mice. A cocktail of two viruses, AAVDJ-FLEX-ChIEF-tdTomato and AAV5-mCherry-Cre, was injected into the dentate gyrus to achieve Cre-specific expression of fast channel rhodopsin ChIEF and PCP-GFP in the granule cells (GC) (**Fig 2G**). A 25 Hz optical stimulation was delivered to the GC layer of dentate gyrus (**Fig 2H**), and transcription from both alleles in individual neurons was imaged for over four hours using two-photon microscopy (**Fig 2I**). In agreement with activity-dependent induction, neurons following optical stimulation showed an immediate early response of *Arc* transcription at 30 min followed by a decline (5.7 ± 1.1% at basal vs 14.6 ± 2.2 % at 30 min) (**Fig 2J**). A subsequent increase in transcribing neurons to 9.7 ± 0.8 % occurred at 150 min and maintained till 180min. This second transcriptional phase primarily consisted of GCs undergoing reactivation and sustained transcription, and a small fraction of delayed *de novo* transcription (**Fig 2K**). Importantly, the time frame of reactivation (120-180 min) was similar to that observed in cultures, suggesting that the temporal regulation of *Arc* gene transcription was conserved in brain tissue. Hence, a better controlled (optical) method for triggering and imaging long-term *Arc* dynamics in cultures and tissue emphasized the generality of the biphasic transcriptional response.

### Reactivation occurs independently from increase in nuclear calcium

Elevated levels of somatic and nuclear calcium have been implicated for induction of IEG transcription (Yap and Greenberg, 2018). Therefore, we investigated the dependency of transcription reactivation on Ca^2+^ activity. First, nuclear Ca^2+^ levels were measured using the indicator NLS-tagged jRGECO1a at different time points after stimulation (**Fig S3A**). An immediate increase in the frequency of Ca^2+^ transients (CaTs) at 10 min was observed, which peaked at 30 min and maintained until 60 min (**Fig S3B**, **C**). This was followed by a significant reduction at 120 min, and negligible activity was observed at 180 min (only 8 out of 40 cells exhibited CaTs). The amplitude of CaTs showed a similar profile: a rapid increase followed by a decrease at 120 min (**Fig S3D**). These findings showed a severe dampening of nuclear CaTs at the two-hour time point when reactivation was evident.

To confirm that reactivation occurred independently of Ca^2+^ rise, neuronal activity was silenced by reapplying TTX to the imaging media at 90 min (**Fig 3A**). Co-imaging of nuclear Ca^2+^ and *Arc* transcription was performed by expressing NLS-jRGECO1a and PCP-GFP in the same neuron (**Fig 3B, C**). Increase in nuclear Ca^2+^ levels and concomitant *Arc* transcription was observed in the same neuron in the IE-phase. After reapplication of TTX (**Fig 3B**, **Movie S3A**), CaTs were abolished. However, reactivation of *Arc* transcription was induced in those same neurons even though nuclear Ca^2+^ levels were undetectable (**Fig 3C**, **Movie S3B**). Despite being lower than TTX-w, a notable 45 ± 2.9 % of neurons displayed reactivation after TTX reapplication (**Fig 3D**), with no significant difference in transcription onset times (154.3 ± 4.2 for TTX-w versus 145.3 ± 6.4 for TTX-w + TTX, **Fig 3E**). Therefore, while the IE-phase was synchronous with elevated Ca^2+^ activity in the neuron, the reactivation onset appeared uncorrelated. This indicated that the later transcriptional phase was governed by a mechanism distinct from the conventional excitation-transcription coupling involving Ca^2+^ rise in neurons.

**Fig 3:**
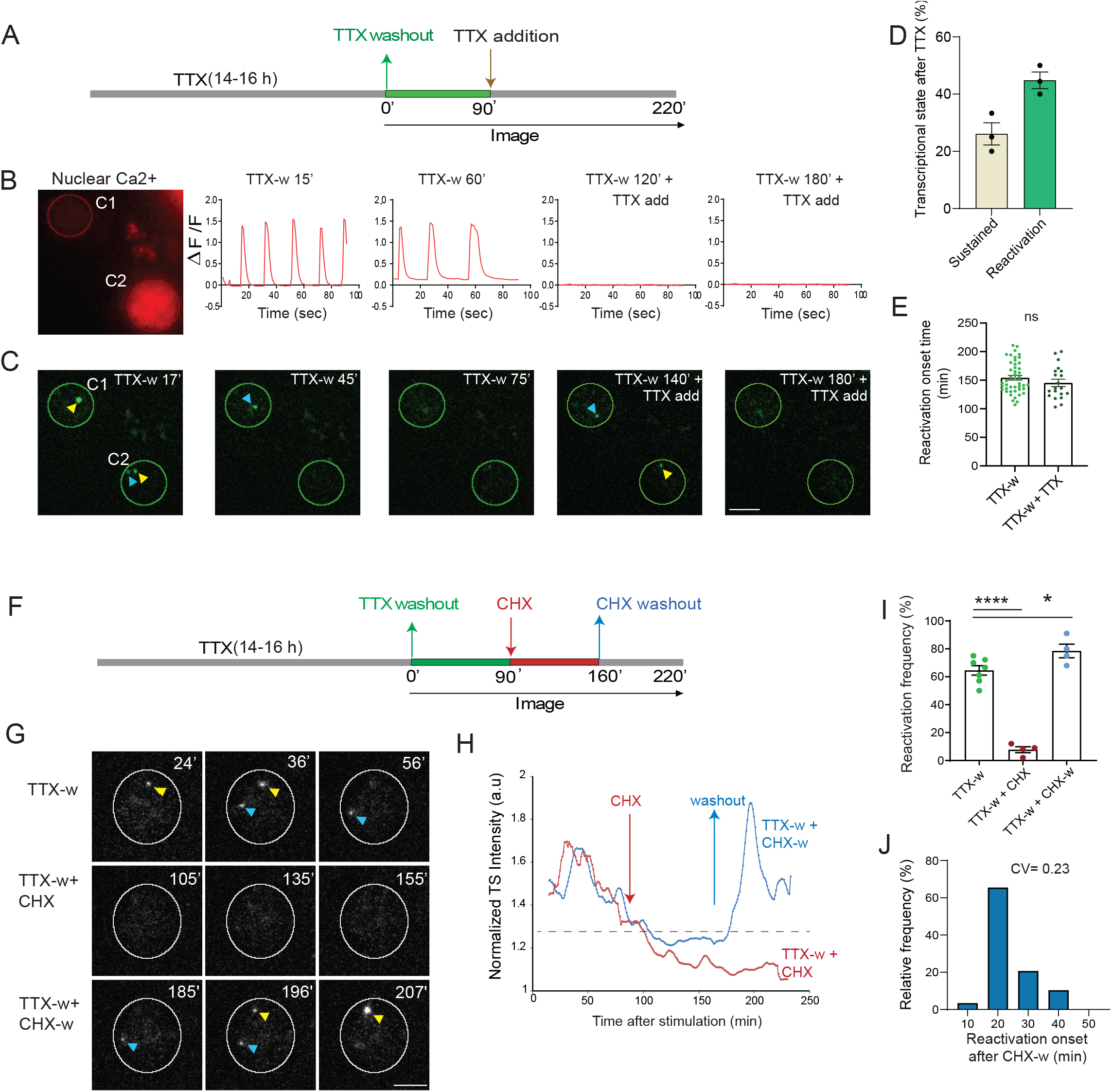
Reactivation of *Arc* transcription is independent of Ca^2+^ rise but requires protein synthesis. (**A**) Schematic of stimulation paradigm. (**B-C**) Neurons co-expressing NLS-jRGECO1a and PCP-GFP were imaged for nuclear Ca^2+^ levels and Arc transcription. Nuclear CaTs triggered by TTX-w ceased after reapplying TTX (**B**). Imaging TS in the same neuron showed reactivation even after TTX addition (**C**). (**D**) Different transcriptional states after TTX reapplication in neurons activated in IE-phase (n = 43 neurons from 3 independent experiments). Delayed *de novo* was not observed. (**E**) Comparison of reactivation onset times (n = 20 neurons for TTX-w + TTX; 45 neurons for TTX-w, p = 0.24). (**F**) Schematic of stimulation paradigm to monitor the effect of protein synthesis. Translation elongation inhibitor (CHX, 50 μg/ml) was added at 90 min (TTX-w + CHX). In another set of experiments, CHX was incubated for 70min and then washed out (TTX-w + CHX-w). (**G**) Representative images showing IE-transcription from both alleles, followed by shutdown maintained with CHX addition. Washout of CHX restored transcription. (**H**) Intensity trace of transcribing alleles from two conditions-CHX addition (TTX-w + CHX), and washout (TTX-w + CHX-w). (**I**) Percentage of reactivation across conditions, each circle represents one experiment (TTX-w vs TTX-w + CHX, **** p < 0.001, TTX-w vs TTX-w + CHX-w, * p = 0.036, one-way ANOVA; n = 21 neurons for TTX-w + CHX, n = 47 neurons for TTX-w + CHX-w, from 4 independent experiments, TTX-w from Fig 1E) (**J**) Frequency distribution of reactivation onset times after CHX washout. CV= coefficient of variation. Error bars indicate SEM. **** denotes p < 0.001, * denotes p < 0.05. Scale bar 5μm.

### Transcriptional reactivation requires new protein synthesis

Classically, IEGs including *Arc,* transcribe rapidly in response to stimulation, without the requirement for new protein synthesis (Greenberg and Ziff, 1984; Saha and Dudek, 2013). We therefore assessed the role of *de novo* protein synthesis for the second transcription cycle. Translation was stalled using cycloheximide (CHX) at 90 min after the initial stimulation (**Fig 3F**). CHX addition severely impacted reactivation, only 7.8 ± 2.1% activated neurons were transcribing two hours later (**Fig 3G-I**). Importantly, removal of CHX resulted in a robust induction of transcription (**Fig 3G-I**) with synchronous initiation times (median = 20 min after washout) (**Fig 3J**, **Movie S4**). To evaluate whether protein synthesis-dependent transcription occurred in tissue at later stages, acute hippocampal slices were briefly depolarized with KCl, and maintained for two hours with or without CHX, and fixed. In another set of slices, CHX was washed out and fixed (**Fig S4A**). Analysis of transcription sites (TS) in fixed slices revealed that neurons transcribing at two hours were significantly reduced with CHX application compared to no CHX, but were rescued upon washout (**Fig S4B, C**). Moreover, transcription from both alleles was also restored (**Fig S4D**). Taken together, these findings indicate that the second phase of transcription requires *de novo* protein synthesis. We postulate that stalling translation elongation maintains a pool of readily translatable mRNA, which leads to a burst of protein synthesis upon CHX washout. These new proteins could then feedback to the nucleus to induce *Arc* gene transcription.

### Arc mRNAs and proteins are maintained in the dendrites over long term after an initial stimulus

Given that at least two cycles of mRNA synthesis occurred, we assessed whether that led to corresponding changes in *Arc* mRNA levels in the dendrites. Time-lapse imaging of single mRNAs showed (**Fig S5A, B**) that the RNA number in dendrites was not constant but displayed fluctuations over time (**Fig S5C**, **Movie S6**). The RNA density plot in dendrites across multiple neurons displayed two phases – first one at 90 min followed by a plateau, and a second peak at 210 min (**Fig S5D**). Since the residence time of the Arc mRNAs in the dendrites was short (average 7.6 ± 1.2 min) (**Fig S5E**), we propose that the increase in RNA density during the second phase was not due to the long-term persistence of mRNAs but resulting from new *Arc* mRNAs populating the dendrites over time.

Additional mRNAs being transported and localized in the dendrites would result in local protein synthesis. To examine whether Arc proteins in the dendrites were replenished over time, a reporter was designed to identify Arc proteins from the different cycles. The reporter driven by an activity-regulated ESARE promoter (Kawashima et al., 2009), contained a HaloTag upstream of the Arc coding sequence (CDS), followed by the 3’UTR comprising the *cis-acting* regulatory elements (**Fig 4A**). Cell permeable Halo-ligand conjugated to JF dyes (Grimm et al., 2017) were used to label the Halo-tagged Arc proteins, similar to an approach used for β-actin (Yoon et al., 2016). Spectrally distinct JF dyes-JF646 and JF549 dyes, detected Arc proteins synthesized from the IE and the second phase respectively (**Fig S5F**). Since inducible Arc translation peaks at two hours (Shepherd and Bear, 2011), labeling with JF646 was performed till 150min post stimulation to detect Arc proteins from the IE-phase. Interestingly, the labeled Arc protein was not evenly distributed, but displayed discrete puncta along the dendrites (**Fig S5G**). A subsequent chase with JF549 for 60 min revealed a second phase of Arc synthesis, where the newly labeled proteins were in close proximity or overlapped with the previous 646 signal. The distances between the brightest JF646 puncta to the nearest JF549 signal were measured and majority (75 %) were within 3.4 μm, indicating the enrichment of Arc proteins from the first and second phases in discrete dendritic domains (“hubs”) (**Fig S5H**). We propose that these hubs potentially represent sites where dendritic Arc proteins accumulate and are maintained over time.

**Fig 4:**
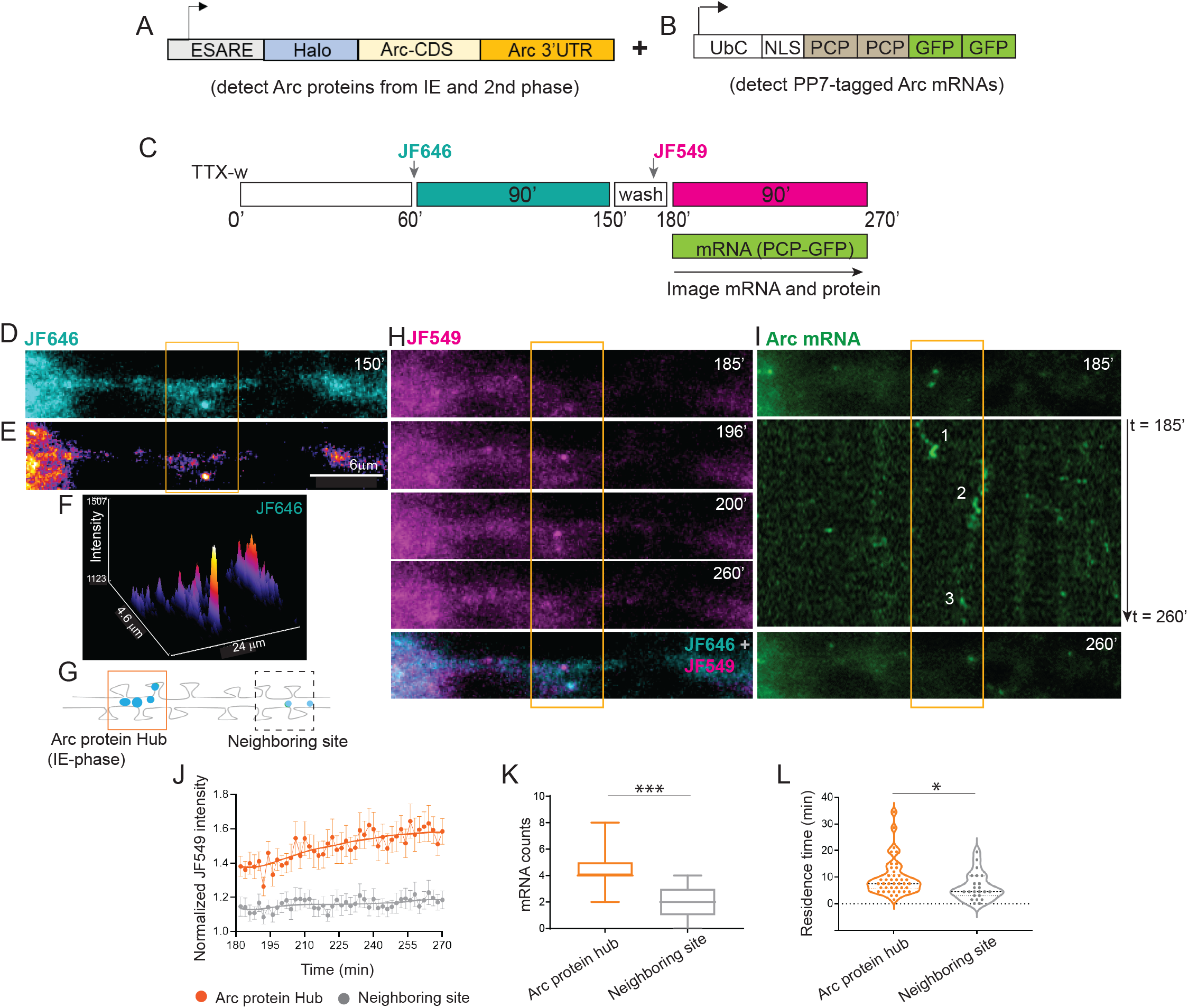
Detection of Arc protein hubs in dendrites and mRNA localization to hubs. (**A**) Schematic of the Halo-Arc protein reporter driven by ESARE (activity-regulated promoter) to detect proteins from IE and second phase. (**B**) PCP-GFP construct to visualize endogenous *Arc* mRNAs. A cocktail of lentiviruses expressing A and B was used to image Arc proteins and mRNAs in the same neuron. (**C**) Schematic of JF646/JF549 labeling timeline after stimulation. Since transcriptional reactivation occurred at 150 min post stimulation, Arc mRNAs and proteins from the second phase were imaged 3 hr onwards. (**D-E**) Representative image of dendritic Arc protein following JF646 labeling during the IE phase of protein synthesis. Differential intensity of JF646 signal shown in **E**. (**F**) Intensity profile of JF646 intensity showed distinct peaks in specific dendritic regions. (**G**) A 6 μm segment around the local maxima in F was used to designate Arc protein hub from IE-stage (orange outline). A region of same dimension (dashed outline) nearby the hub was marked as neighboring site. (**H**) Labeling with JF549 shows Arc proteins synthesized in the second phase. Bottom panel shows a merged image of JF549 with JF646, indicating close proximity of both puncta. (**I**) Localization of endogenous *Arc* mRNAs in the same dendrite. Middle panel shows a kymograph, where localized mRNAs are numbered. (**J**) Normalized intensity trace of new Arc protein (JF549 signal) over time in hub versus neighboring site as defined in G. (**K**) Comparison of Arc mRNA counts populating the hubs versus neighboring sites (*** p = 0.002, Wilcoxon signed rank test). (**L**) Comparison of the residence times of Arc mRNAs in the hubs wrt neighboring sites (* p = 0.029). n =12 neurons from 3 independent experiments (J, K). n = 43 mRNAs for hub versus n = 23 mRNAs for neighboring site (L). Scale bar is 6 microns. *** denotes p < 0.005, * denotes p < 0.05. Error bars represent SEM. Scale bar is 6 microns.

### Arc protein hubs are consolidated over time and serve as landing sites for Arc mRNAs from the second transcription cycle

If indeed the hubs were selectively consolidated compared to other dendritic regions, then repeated enrichment of *Arc* mRNAs and proteins should occur in these hubs. To test this possibility, the localization kinetics of mRNAs and proteins from the second phase was assessed relative to the hubs with high temporal resolution. A three-color real-time imaging approach was developed, where neurons from the Arc-PBS mouse were infected with two lentiviruses: one expressing PCP-GFP and the other expressing the Arc protein reporter (**Fig 4A, B**). The timeline of labeling and imaging has been depicted in the scheme (**Fig 4C**). A detailed analysis of JF646 signal revealed that Arc protein from the IE phase was spatially concentrated (**Fig 4D, E)**. The local maxima of JF646 puncta intensity were used to mark a segment around the centroid of the peak (3 μm on either side, based on Fig 5H) to designate the Arc hub from the IE phase (**Fig 4F, G)**. Time lapse imaging with JF549 showed that Arc proteins translated in the second phase congregated in the region of the existing hub (**Fig 4H**). However, these JF549 puncta were not long-lasting, suggesting possible degradation of the Halo-Arc protein. Importantly, the endogenous *Arc* mRNAs in the same dendrite also exhibited localization at or in the vicinity of the existing hubs, as shown by the kymographs (**Fig 4I**).

**Fig 5:**
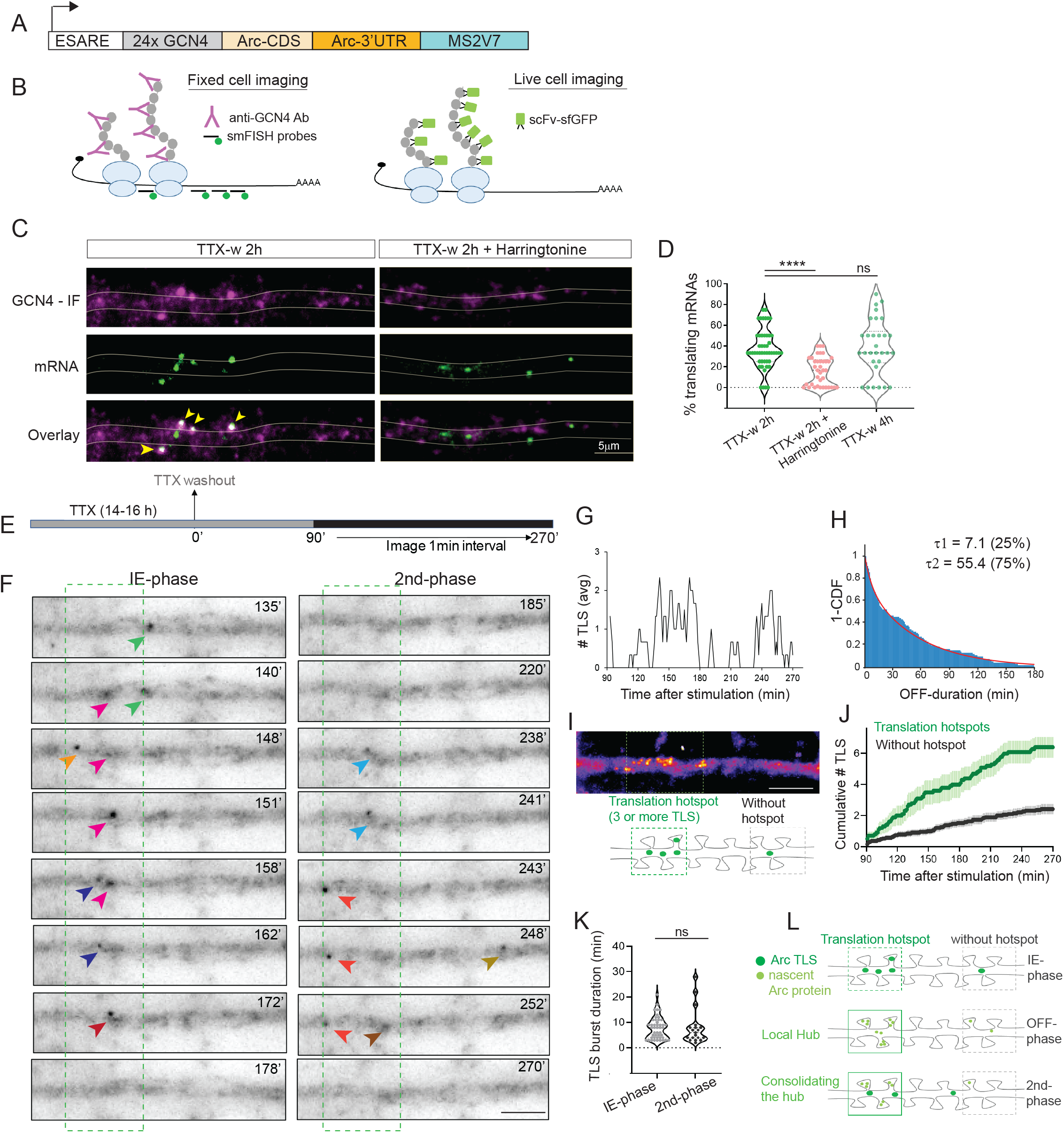
Long-term imaging of Arc translation reveals hotspots and biphasic dynamics. (**A**) Schematic of the Suntag-Arc translation reporter. The reporter is driven by ESARE (activity-regulated promoter) and contains the 24X GCN4 epitopes of Suntag upstream of the Arc CDS and followed by the 3′ UTR of Arc, and stem loops MS2V7. (**B**) The translating mRNAs are detected by fixed-cell imaging with antibodies against GCN4 and by smFISH against GCN4 and stem loop sequence. In live cells, translation sites (TLS) are detected by binding of the single chain antibody against GCN4 (scFV) fused to superfolder GFP (sfGFP). (**C**) Images of dendrites showing both nascent peptides and mRNAs in stimulated and after inhibition with Harringtonine. Co-localization of IF-smFISH spots indicate translating mRNAs (yellow arrows). (**D**) Comparison of translating mRNAs after stimulation and translation inhibition (TTX-w 2h vs TTX-w 2h + Harringtonine, **** p < 0.001, TTXw 2h vs TTXw 4h, p = 0.23, one-way ANOVA; n = 49 dendrites for TTX-w 2h, n = 34 dendrites for TTX-w 4h, n = 37 dendrites for TTX-w + Harringtonine from 2 independent experiments). (**E**) Stimulation and imaging paradigm to capture long-term dynamics of Arc TLS. (**F**) Images from a dendrite showing appearance of TLS (arrows). Different colors represent different TLS. Time was binned to represent IE (90-180 min) and second (181-270 min) phase. Green dotted box indicates the ROI with TLS clustering, representing hotspots along the dendrite. (**G**) TLS counts (rolling average 3) show biphasic dendritic translation. (**H**) Inverse cumulative distribution function of OFF-periods fitted to a 2-component exponential, yielding two time constants τ1, τ2, and relative percentage of events indicated (n = 90 events). (**I**) A time-projected image showing spatially clustered Suntag signal forming a translation hotspot. Lower panel shows schematic where at least 3 TLS tracks in 8μm segment was used as the inclusion criterion for a hotspot. (**J**) Cumulative number of TLS in hotspots versus regions without hotspots (n = 20 dendrites, 3 independent experiments). (**K**) Duration of translation bursts during the IE and second phase (n= 35 events in IE, n=22 events in second phase). (**L**) Proposed model of emergence and maintenance of Arc protein hubs by local translation in the hotspots. *Arc* translation in specific hotspots during IE-phase promote rapid increase in nascent protein density to form the “hubs”. During the OFF-phase, translation is low possibly due to limited mRNA availability. A second transcriptional cycle supplies mRNAs, which preferentially visit the hubs to undergo a second phase of translation, and consolidate the hub compared to the neighboring dendritic regions. Error bars indicate SEM. **** denotes p < 0.001, * denotes p < 0.05, ns for p > 0.05.

Quantification of the Arc protein from the second phase showed elevated levels specifically in the hub over time compared to a neighboring dendritic segment (**Fig 4J**). The peak of new Arc enrichment in the hub occurred at 252 mins after a latency period, supporting the idea that these hubs are being reinforced by cycles of new protein synthesis. To establish that subsequent Arc protein synthesis and consolidation of the hub is driven by newly transcribed *Arc* mRNAs, DRB (a transcription blocker) was added after the IE-phase was over. Indeed, inhibition of the second transcriptional cycle prevented the Arc protein enrichment in the hub over time (**Fig S6**).

Notably, there was a spatial correlation between Arc mRNA localization in the second phase with the protein hubs from the IE phase; a two-fold higher enrichment of Arc mRNAs in the initial hub versus the neighboring site was observed (**Fig 4K**). Once localized, the mRNAs persisted at these hubs as calculated from their residence times (11.9 ± 1.9 min in hub versus 6.3 ± 1.1 min in neighboring site) (**Fig 4L**). These results revealed distinct spatial and temporal features of dendritic *Arc* mRNAs and their cognate proteins: i) a biphasic regulation of Arc protein synthesis separated by 2 hours, and ii) the emergence of local hubs of Arc proteins, which are reinforced over time by preferential localization of *Arc* mRNA from later cycles of transcription.

### Long-term translation imaging reveals cycles of local Arc translation in dendrites

To determine whether local translation at certain dendritic hotspots generated the Arc protein hubs, and the temporal regulation of the process, translation was imaged in real time with multimerized epitopes (“Suntag”) on the Arc protein and detected with a genetically encoded single chain antibody (scFV) fused to a fluorescent protein (Tanenbaum et al., 2014; Wang et al., 2016; Wu et al., 2016). The Suntag-Arc reporter contained 24x repeats of the epitope in the N-terminus of Arc CDS with Arc mRNA 5′ and 3′UTRs for translation regulation (**Fig 5A**). Fusing the 24x Suntag did not alter the localization of Arc protein (**Fig S7A**). The ability of the Suntag-Arc construct to report translating Arc mRNAs was tested by performing single molecule FISH (smFISH) for the mRNA and immunofluorescence (IF) against the Suntag protein in fixed neurons **Fig 5B**, left panel). Co-localization of smFISH spots with bright Suntag IF-signal showed mRNAs undergoing translation at two hours post stimulation. This was significantly reduced upon addition of Harringtonine (inhibitor of translation initiation) (37.2 ± 2.4 % TTXw 2h vs 16 ± 2.3 % Harringtonine, **Fig 5C, D**). Notably, the translating Arc mRNAs were maintained at 4 hour post stimulation, indicating that the mRNAs from early and late phases were translationally competent.

To investigate the spatio-temporal dynamics of local translation over time, Suntag-Arc reporter and scFV-GFP (**Fig 5B**, right panel) were co-expressed and imaged for several hours after stimulation (**Fig 5E**). Discrete foci, much brighter than the faster diffusing proteins were detected in the dendrites (**Fig 5F, Movie S5**), indicative of nascent sites of translation (TLS)(Wu et al., 2016). Tracking these TLS revealed that their numbers were not constant over time but displayed cycles of increase and decay (**Fig 5G**). Distribution of the OFF periods where no TLS are detected revealed a long t2 component of 55.4 ± 8.3 min (**Fig 5H**). These measurements closely mimicked the OFF periods of transcription (∼1hr), suggesting a temporal coordination of transcription and translation cycles. Spatial analysis of TLS distribution showed that they were not homogeneous but clustered along the dendrites (**Fig 5I**). This is representative of translation hotspots which are potential sites for increased translation efficiency (Job and Eberwine, 2001a). Accordingly, the appearance of multiple TLS (3 or more) in an 8μm dendritic segment within 2-hour post stimulation was used as a criterion to define a translation hotspot. A cumulative increase of Arc TLS continued in these hotspots after IE-phase at a considerably higher rate than in a region without a hotspot (**Fig 5J**), closely resembling the patterns of spatially selective Arc protein enrichment (Fig 4). The abundance of nascent Arc proteins within the translation hotspots possibly facilitates the formation of local hubs.

The temporal kinetics of Arc translation in these hotspots was obtained by binning the graph in Fig 5I into two 90-minute bins, congruent with the two phases of Arc protein detection in Fig 4. The cumulative TLS counts revealed increased magnitude, indicative of more translation events during the IE phase compared to the second phase (**Fig S7B, C**); mimicking the mRNA output measurements during transcriptional reactivation (Fig S2). Furthermore, tracking the position and the integrated intensity of individual TLS from the two phases demonstrated that translation occurred in bursts lasting an average of ∼8 min/burst, irrespective of the phase (**Fig 5K**). Hence, intermittent bursts of protein synthesis at local translation hotspots could be a mechanism by which nascent Arc proteins are organized in the hubs and maintained over time (**Fig 5L**).

## Discussion

In this study, the long-term dynamics of the *Arc* gene from transcription to translation in live neurons were followed using high-resolution imaging. This identified a novel temporal regulation of *Arc*, an IEG with critical roles in long term memory. Cycles of transcription and local translation demonstrated that intermittent phases of mRNA and protein synthesis maintain dendritic Arc protein levels over time. The protein is organized as local hubs, possibly because of local translation in certain hotspots and consolidated over time by subsequent translation at the same hotspot. These dynamics beyond the IE-phase provide a potential molecular mechanism by which a transient synaptic protein like Arc impacts long term spine remodeling and in turn memory storage.

The molecular events that occur inside the neuron during memory formation including the stabilization of synaptic contacts, have not been elucidated completely. It has been apparent that mRNA localization to activated spines plays a role in this process, and this leads to localized translation of specific proteins necessary for the structural and functional integrity of the post- synaptic structures (Costa-Mattioli et al., 2009; Das et al., 2021; Holt et al., 2019; Sutton and Schuman, 2006; Swanger and Bassell, 2011; Wang et al., 2009). To support memory consolidation, the activity-driven changes in the synapses must be sustained (Kelleher et al., 2004). For a structural protein such as β-actin, its mRNA is constitutively expressed, abundant, long-lived and can persist at, or revisit sites of activity to translate and promote stability of the cytoskeletal architecture within the postsynaptic region (Buxbaum et al., 2015; Yoon et al., 2016). The plasticity protein, Arc, is synthesized from mRNAs transcribed upon activity, after they traffic to sites of activity to be translated locally (Na et al., 2016; Steward et al., 2014; Steward et al., 1998). However, in contrast to actin, both Arc mRNAs and proteins are transient and degrade within a couple of hours (Farris et al., 2014; Rao et al., 2006). For synaptic structures and physiology to be maintained, Arc must be persistently concentrated there, and this likely occurs via a second mechanism: cycles of transcription and translation in response to an initial stimulation. Since Arc is important for long term memory (Guzowski et al., 2000; Plath et al., 2006; Shepherd and Bear, 2011), this cyclical regulation that begets additional rounds of transcription and protein synthesis creates a constant feedback loop that presumably supports memory consolidation.

Sustained levels of *Arc* mRNAs and proteins have been observed in cultured neurons (Kawashima et al., 2009), in different areas of the hippocampus after spatial learning tasks and during long term memory (Igaz et al., 2002; Nakayama et al., 2015; Ramirez-Amaya et al., 2013). However, it has been challenging to distinguish whether one or more transcription events were occurring in the same or different neurons. Using the Arc-PBS mouse and the new Arc translation reporters, we characterized the different phases of gene expression with unprecedented temporal resolution. Reactivation of transcription in the same neuron predominantly accounted for the transcriptional events beyond the IE phase in both cultures and in tissue (Figs 1, 2), irrespective of the stimulation condition, suggesting a conserved intrinsic feature of the *Arc* gene. Since, additional depolarization is not a requirement (Fig 3), transcriptional reactivation may occur by i) signaling cascades activated during IE-phase that remain sustained over long periods, and/or, ii) transcription factors which are synthesized in the IE phase and do not require Ca^2+^-mediated activation. The dependency of reactivation on protein synthesis (Fig 3) provided evidence favoring the latter possibility. Indeed, several TFs capable of regulating *Arc* expression directly or indirectly are rapidly translated IEGs, such as zif268 (Li et al., 2005; Penke et al., 2011), Npas4 (Sun and Lin, 2016). Therefore, the molecular regulation of the two transcriptional phases is distinct due to their differential requirement of Ca^2+^ and *de novo* protein synthesis.

The phenomenon of reactivation is not just restricted to transcription; local translation of *Arc* mRNAs in the dendrites also occurs with a periodicity of ∼2 hours. Reactivation could have several benefits. First, it allows production of short bursts of mRNAs and proteins without saturating the system. This is important for Arc, the levels of which need to be strictly regulated for cognitive flexibility (Wall et al., 2018). Second, reactivation maintains the RNA levels and efficiently replenishes the Arc hubs over long term. Finally, reactivation of Arc expression in a subset of neurons could favor their recruitment to a neuronal assembly supporting the memory trace (Asok et al., 2019). Therefore, the reactivation signature could potentially identify neurons involved in circuit strengthening during memory consolidation.

The formation and maintenance of the local Arc hubs by periodic translation in hotspots highlight that the consolidation of Arc protein along the dendrites is spatially selective. These hotspots may either indicate regions of elevated synaptic activity where remodeling of spines is persisting, and/or regions of increased ribosome density (Sun et al., 2021) corresponding to high rates of translation (Job and Eberwine, 2001a)(reviewed in (Das et al., 2021; Job and Eberwine, 2001b). Such hotspots of ribosomes and nascent proteins are often correlated with increased spine density, and prevalent in both local stimulation as well as global paradigms (Sun et al., 2021). In our case, translation in the hotspots allows the nascent Arc protein to accumulate locally to form the hubs. These local high concentrations of Arc could also facilitate the generation of Arc capsids for intercellular communication (Pastuzyn et al., 2018).

In the last decades, studies have independently focused on the contribution of transcription and local translation to long-term memory (Alberini and Kandel, 2014; Asok et al., 2019; Costa-Mattioli et al., 2009; Sutton and Schuman, 2006). However, little is known about how these two important cellular processes converge to regulate memory consolidation. We have shown that coupling between transcription and translation is maintained for the subsequent cycles several hours after stimulation, consistent with the timing of stabilization of activity-driven changes. Cyclical gene expression may be a feature of strictly timed genes, such as seen for developmental genes (Hendriks et al., 2014; Kim et al., 2013), and future studies characterizing the persistence of Arc cycles during memory stabilization are needed. The long-term dynamics of *Arc* gene expression may serve as a template for studying other IEGs involved in memory.

Determining the ubiquitous nature of these transcription-translation cycles, and whether they are synchronized or sequential for specific IEGs is critical to advance our knowledge of brain functions. Improvements in technologies to image mRNAs and proteins *in vivo* will pave the way towards understanding the transcription-translation coupling after a learning task, and whether perturbations of these cycles affect consolidation and persistence of memory.

## Acknowledgments

We are grateful to Xiuhua Meng and Melissa Lopez-Jones for technical assistance, and Chiso Nwokafor for animal maintenance. We thank all members of Robert H. Singer and Pablo E. Castillo laboratories for insightful discussions. We are grateful to Luke Lavis from Janelia Research Campus for providing the JF646 and JF549 dyes. We would like to thank Dong-woo Hwang and Weihan Li for critical reading of the manuscript.

The work is supported by:

National Institutes of Health grant R21MH122961 (SD) National Institutes of Health grant R01NS083085 (RHS)

National Institutes of Health grant R01MH125772, MH116673, NS115543, NS113600 Ruth L. Kirschstein NRSA Fellowship MH109267 (PJL)

## Author contributions

S.D. conceptualized the project, and with R.H.S. designed the research. S.D. performed real-time imaging of transcription, translation in cultures and analyzed all experimental data. P.J.L. performed stereotaxic surgeries, and two-photon microscopy of transcription in slices. All authors discussed the results. S.D. and R.H.S wrote the first manuscript, which was edited by P.E.C. and P.J.L.

## Data and materials availability

All data, code, and materials used in the analysis will be made available upon request.

## Methods

### Constructs and Viruses

All lentiviral constructs were cloned into the phage-ubc-RIG lentiviral vector. For Arc-translation construct, 24X Suntag or GCN4 repeats were PCR-amplified from the SINAPs construct (Wu et al., 2016). The promoter was the minimal ESARE promoter previously described (Kawashima et al., 2009) and synthesized as a gene block along with the 5’UTR sequences. The Arc coding sequence was synthesized based on the sequence (NM_001276684.1), and the 3’UTR sequence was a kind gift from Xiaowei Zhuang (Wang et al., 2016). The 3’UTR sequence was inserted before the woodchuck hepatitis virus posttranscriptional regulatory element (WPRE) in the lentiviral backbone.

The Halo-Arc protein reporter was designed based off the Halo-Actin-reporter (Yoon et al., 2016), where the β-actin coding sequence and β-actin 3′UTR were replaced by the Arc CDS and Arc 3′UTR respectively. For PBS coat protein, PCP, we used the synonymously transformed version fused to GFP stdPCP-stdGFP (Das et al., 2018).

The construct for red-shifted nuclear calcium indicators, NLS-jRGECO1a were cloned into the p323 backbone. The sequence for jRGECO1a was obtained from Addgene #61563, and PCR amplified with primers to add NLS to the N-terminus. ChR2-mCherry construct was obtained from Addgene (#20938) and subcloned into p323 backbone. Lentiviral particles were produced by transfecting the expression vector with accessory plasmids, ENV, REV, VSVG and GAG in HEK 293T cells. Collected lentiviral particles were purified with lenti-X concentrator (Clontech, Mountain View, CA).

### Animals

Two animal strains were used-the Arc PBS-KI line and the Arc^P/P^ x PCP-GFP line. The Arc-PBS line has been previously published and have been maintained at homozygosity. The genotyping primers and the conditions have been described in an earlier study (Das et al., 2018). The PCP-GFP mice were generated by introducing a neocassette containing CAG-stop/flox-NLS-PCP-GFP in the ROSA 26 locus, where PCP-GFP expression is Cre-inducible. The Arc-PBS mouse (Arc ^P/P^) were crossed with the PCP-GFP animals to generate the The Arc^P/P^ x PCP-GFP line are genotyped using the following PCRs for these primer sets: R26 wt forward primer (5′-CCAAAGTCGCTCTGAGTTGT-3′), and reverse primers, R26 wt (5′- CCAGGTTAGCCTTTAAGCCT-3′), and CMV R1 (5′-CGGGCCATTTACCGTAAGTT-3′), yielding a 250 bp product for the WT allele, and a 329 bp product for the PCP-GFP respectively. Arc PBS were genotyped to All animals are being maintained at homozygosity with routine genotyping at Transnetyx. All animals are being maintained according to IUCAC guidelines.

### Stereotaxic Surgery and Hippocampal slice preparation

Arc^P/P^ x PCP-GFP mice at postnatal day 27 (P27) were anesthetized with oxygen-isoflurane flowing at 1.5 ml/min and positioned into a Kopf stereotaxic instrument. A beveled Hamilton syringe injected 1-1.5 μl of 1:2 mix of AAV5-CaMKII-mcherry-Cre/AAV-DJ-FLEX-ChIEF-tdTomato virus at coordinates targeting the dentate gyrus (−2.1 mm A/P, 1.7 mm M/L, 2.5 mm D/V). Animals were sacrificed 5-weeks post-surgery using anesthesia (4% Isoflurane) followed by decapitation, and slices were prepared for two-photon microscopy. No differentiation of sex was done. Briefly, the animals were perfused with 20 mL of cold N-Methyl-D-glucamine (NMDG) solution containing in (mM): 93 NMDG, 2.5 KCl, 1.25 NaH2PO4, 30 NaHCO3, 20 HEPES, 25 glucose, 5 sodium ascorbate, 2 Thiourea, 3 sodium pyruvate, 10 MgCl2, 0.5 CaCl2, maintained at pH 7.35. The hippocampi were isolated and cut (300 µm thick) using a VT1200s microslicer in cold NMDG solution. Acute hippocampal slices were placed in a chamber with artificial cerebral spinal fluid solution (ACSF) solution composed of 124 NaCl, 2.5 KCl, 26 NaHCO3, 1 NaH2PO4, 2.5 CaCl2, 1.3 MgSO4 and 10 glucose (in mM), and incubated in a 33-34°C water bath. All solutions were equilibrated with 95% O2 and 5% CO2 (pH 7.4). Post-sectioning, acute slices were allowed to recover at room temperature for at least 45 min prior to experiments.

### 2-photon imaging in slices and optical stimulation

An Ultima 2P laser scanning microscope (Bruker Corp.) equipped with an Insight Deep See laser (Spectra-Physics) tuned to 910-930 nm was used to image *Arc* transcription in dentate granule cells (GCs) with 512 x 512 pixel resolution using 4 mW laser power measured at the 60X objective (Nikon, 1.0 NA). GCs expressing the PCP-GFP coat protein were imaged at 1X magnification to detect at least one transcribing neuron, which was then chosen as the region of interest (ROI) for 2X magnification. A Z-stack of 25 µm thickness with 0.5 µm steps was taken to assess baseline Arc transcription signals before optical stimulation.

Acute slices showing optimal ChIEF-tdTomato reporter expression (at least 75% of DG was fluorescent) were selected for optical stimulation and imaging. The Invitro Ultima 2P microscope (Bruker Corp.) contains a Coherent 473 nm laser path that delivered optical stimulation of 25 pulses at 25 Hz repeated 20 times every 5 s (8 mW, 2-4 ms pulse duration). The stimulation area was specifically defined using customized Mark Point software (Bruker Corp.) and was empirically determined based on at least one transcribing neuron in the field of view as described above. After stimulation, Z-stack images of 25 µm thickness with 0.5 µm steps were acquired every 15 min for 4-5 hour.

### Depolarization of Slices and imaging of fixed slices

Arc-PBS X PCP-GFP animals injected with AAV5-CaMKII-mcherry, were sacrificed 3-weeks post injection and acute hippocampal slices were prepared. The slices were briefly depolarized with 90 mM KCl for 3 mins and returned to ACSF at room temperature (RT). After 90 mins, slices were grouped into three treatment conditions: first group in ACSF, second group was incubated with CHX (100mg/ml) for 1 hour, and in the third group, CHX was added for 1 hour followed by washout. The first two groups were fixed at 2.5 hours, and the group with CHX-washout was fixed 45 min later to allow complete washout. Fixation was done with 4% PFA in PBS overnight, and washed thrice with PBS, and then mounted onto slides with Prolong Diamond (Invitrogen). Imaging was performed on a wide-field fluorescence microscope built around an IX-81 stand (Olympus) and illumination was with 488nm laser, captured on EMCCD camera (Andor, iXon3 DU-897E-CS0-#BV). 300nm z-stacks were acquired, and max-projected to obtain the images of GCs.

### Primary Hippocampal Neurons and Stimulation paradigms

Mouse hippocampi were isolated from Arc PBS or Arc-PBS x PCP-GFP animals at post-natal day 0 or 1 (P0/P1). The tissue was digested in 0.25% tryspin for 15 min at 37 °C, triturated and plated onto Poly-D-lysine (Sigma) coated glass bottom Mattek dishes at a density of 75,000 cells/dish for live imaging and 60,000 cells/dish for fixed cell imaging. Primary neurons were cultured in Neurobasal A media supplemented with B-27, GlutaMax and primocin (InvivoGen). Viral transduction was usually done at or after DIV 7, and neurons were imaged between DIV 16-19.

The paradigm for TTX-washout was used as described before (Das et al., 2018; Saha et al., 2011). Neurons were treated overnight (14-16 hours) with TTX (2μM), followed by washes, and fresh Hibernate A media (Brainbits) was added. In another set of experiments, a chemical LTP paradigm was used (Donlin-Asp et al., 2021). Briefly, neuronal cultures were incubated with 50 μM APV (Tocris) for 12 hours, followed by induction with 200 μM glycine (Sigma) and 100 μM picrotoxin (Tocris) in Mg^2+^-free Hibernate A media for 5 min. Cells were then washed twice and returned to Hibernate A media with calcium and magnesium.

### Imaging in cultured hippocampal neurons

Hippocampal neurons were imaged in Hibernate A media (Brainbits) at 35°C in a humidified chamber. Time-lapse imaging of transcription and translation was performed on a fluorescence microscope built around an IX-81 stand (Olympus) as described before (Das et al., 2018; Wu et al., 2016). The following lasers were used for illumination: 491 nm laser (Calypso-25, Cobolt, San Jose, CA), 561 nm line (LASOS-561-50, Lasertechnik GmbH, Germany) and a 640 nm line (CUBE 640-40C, Coherent Inc, Santa Clara, CA) were combined, expanded and delivered through the back port. The power for all lasers were controlled by an acousto-optic tunable filter (AOTF) (AOTFnC-400.650-TN, AA Opto-electronic).

The lasers were reflected by a four-band excitation dichroic mirror (Di01-R405/488/561/635, Semrock) to a 150x 1.45 N.A. oil immersion objective (Olympus). For transcription imaging, a 60x 1.40 NA oil immersion objective was used. The fluorescence collected by the same objective, were recorded on an EMCCD camera (Andor, iXon3 DU-897E-CS0-#BV). The emission filters (FF01-525/50 for green and FF01-605/64 (Semrock) for red respectively) were mounted on a motorized filter wheel (FW-1000, Applied Scientific Instrumentation) for fast switching between wavelengths. For stimulation of single ChR2-expressing neurons, a size-adjustable pinhole was used in the excitation light path to restrict the illumination to an area of approximately 10 μm in diameter and prevent cross-activation of neighboring cells.

The microscope is equipped with an automated XY-stage (ASI, MS2000-XY) and a piezo-Z stage (ASI) for fast z-stack acquisition. The microscope was controlled and imaging performed on the Metamorph platform. The ChR2 stimulation paradigm was automated with custom journals in Metamorph. Briefly, the 10 μm illumination spot was recorded and the ChR2 expressing neuron was moved to that area, and 5 images were taken. Next, 2 trains of stimulation of that neuron was done using 491nm laser at 20Hz for 20 times at power density (7 mW/mm^2^), and switched back to wide-field illumination without the pinhole in the light path. For most experiments, a total of 11 z-stacks with 400 nm distance between stacks were acquired. Arc mRNAs and translation sites were imaged as z-stacks with 300 nm step size. The stacks were z-projected and used for analysis. For nuclear calcium, imaging was performed at single z-plane with 50ms exposure times at 1Hz acquisition rate.

### Single molecule fluorescence in situ hybridization (smFISH) and immunofluorescence (IF)

Hippocampal neurons plated at 60,000 cells/Mattek dish were transduced with Suntag-Arc translation reporter for 10 days were stimulated with TTX-washout paradigm at DIV 19 and fixed. The neurons were then permeabilized and smFISH-IF was performed according to the protocol described in (Eliscovich et al., 2017). Briefly, 100nM probes against 24X Suntag sequence and primary antibodies against GCN4 epitopes (Clone C11L34, Ab00436-1.4, Absolute antibody, Wilton, UK) was used in hybridization buffer for 3 hours at 37°C. After washes, the cells were incubated with Alexa Fluor 647 secondary antibody (Life Technologies) and mounted using ProLong diamond antifade reagent with DAPI (Life Technologies). Images were taken in a custom up-right widefield Olympus BX-63 microscope equipped with a Lumencor SOLA-Light engine, ORCA-R2 Digital CCD camera (Hamamatsu), SuperApochromatic 60x/1.35 NA Olympus objective (UPLSAPO60XO) and zero pixel shift filter sets: DAPI-5060C-Zero, Cy3-4040C-Zero and Cy5-4040C-Zero (Semrock). Image pixel size: XY, 107.5 nm; Z-steps, 200 nm. The sequences for FISH probes have been described in (Wu et al., 2016).

### Image Analysis

#### Fixed cell analysis

smFISH - IF images were analyzed using FISH Quant (Mueller et al., 2013). Briefly, the FISH spots in the dendrites were filtered and fit to a 3D Gaussian to determine the coordinates of the mRNAs in Cy3 channel. The intensity and the width of the 3D Gaussian were thresholded to exclude non-specific and autofluorescent particles. Similarly, independent analysis of Suntag spots were performed using FISH-Quant. Co-localization analysis was done using by finding mRNAs which had Suntag signal within 300 nm distance.

#### Live-cell imaging data analysis

Transcription site analysis: Time-lapse images of transcription were obtained after maximum intensity projection of the z-series. The transcription sites (TS) in time lapse images were tracked and their fluorescence intensities were quantified with custom programs written in Matlab (Mathworks). The intensity of TS was normalized to the diffusive PCP-GFP signal in the nucleus. Intensity traces were subjected to a rolling average of 3 frames to remove fast fluctuations in fluorescence signal. A value ≥ 1.4 was considered ON-state of the gene (Das et al., 2018). Values below 1.4 were considered OFF-state. An OFF-period of at least 30 mins between two transcriptional bursts, when the second burst was after 90 min post stimulation was used as criterion to designate reactivation. If any of the alleles was transcriptionally active beyond the IE phase and continued till 100 min post stimulation, then the transcriptional state was considered sustained. Induction of de novo transcription after 100 min of stimulation is marked as an delayed (*de novo*) event. To determine the transcriptional bursting parameters in each cycle, the images were binned into two time segments: IE (15-75 min) and reactivation phase (105-200 min). Each phase can be composed of multiple transcriptional bursts (Larsson et al., 2019). The duration of the ON-state, and the area under each burst were quantified and the sum of all the bursts in each phase have been represented.

##### Translation site analysis

To track translation sites (TLS) in neurons, we chose dendritic segments which are >30 μm from soma or in secondary dendrites. Presence of at least one TLS at the start of imaging was used as a criterion for choosing a particular dendrite. Semi-automated tracking of TLS was performed using Trackmate and custom-built program on MATLAB, with a gap of 2 frames allowed to be treated as the same site. TLS which lasted for at least 3 frames were used for analysis. The intensity of TLS was normalized to the diffusive scFv-sfGFP signal in the dendrites. A threshold of 1.5 was used as a cutoff to be considered as a translating spot.

The gradual increase in the intensity of the translation spots indicates more nascent peptide synthesis and therefore deemed to be translating mRNA. The time between the active translation bursts across multiple TLS represented the OFF-durations, which were plotted as an inverse cumulative distribution function (1-CDF) and fitted to a 2-component exponential fit (goodness of fit test performed in MATLAB).

##### Single mRNA analysis

Single particle tracking of mRNAs was performed using Trackmate plugin on Fiji. The dendrites were straightened before analysis. Tracks shorter than 3 frames were not considered. mRNA counts were normalized to the length of the dendrites to determine RNA density. The track lengths of both stationary and moving mRNAs were used to calculate the residence times of mRNAs in dendrites. For Figure 5, kymographs were plotted for straightened dendrites. The number of mRNAs which last for ≥2 frames (3 min) were considered for mRNA counts in the pre-existing versus the neighboring sites.

##### Pulse-chase assay

The puncta from JF646 and JF549 channels were localized with Analyze particles plugin on Image J. The puncta were diffraction limited spots, which could be fitted to a 2D-Gaussian. High intensity diffusible 646 signal was not considered for analysis. The coordinates were noted. The distances were calculated from the brightest JF646 puncta using the nearest neighbor analysis. JF646 puncta whose brightness were above 20% above background fluorescence (diffusive signal in dendrites) was used for analysis. From the frequency distributions of the distances, 75 percentile corresponding to 3.4 μm was used as the cutoff to designate protein enrichment in space. Accordingly, we used a segment of 6 μm length (3 μm on either side of the centroid of the brightest 646 puncta) to define the Arc protein hub from IE-phase.

##### Nuclear Ca^2+^ imaging analysis

Images were acquired at 1Hz frequency at single planes. The time series were used, and the ROIs for the nuclei were detected in a semi-automated manner in Fiji. The same cells were imaged over time and the same ROIs with required correction for x-y drift were used to quantify fluorescence. The average value from the first 3 frames was treated as baseline fluorescence (F), and the following change in fluorescence (ΔF) was measured. The values in the traces are represented as ΔF/F. Traces were fitted to a peak-fitting algorithm and the maximum likelihood analysis was performed for peak assignment. The frequency and amplitude of the peaks were calculated.

#### Transcription imaging in slices

Images of GC nuclei were filtered, and then semi-automated detection of transcription sites was performed based on fitting to a 2D-Gaussian and ROIs of 28-32 pixels was used for each transcribing allele. The intensity of each transcription site was normalized by the background signal of the nucleus using the same ROI dimension. The normalized intensity value of 1 represents that no transcription sites are detected. An intensity threshold of 10% change (i.e. values 1.1 or higher) was used to designate a transcription site. The same transcription site was followed over time to measure the change in normalized intensity values, and below 1.1 was considered as transcriptional shutdown. Quantification of total transcribing cells were performed as follows:

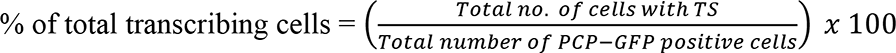

### Statistics

One-way ANOVA (Dunett’s multiple comparison) was used to determine statistical significance for more comparison between more than two groups. Student’s t-test determined statistical significance for all other experimental conditions. Paired t-test was performed when comparing the same neuron or the same allele from the IE and reactivation phases. Normality test for all conditions was performed using the Shapiro-Wilk test. Wilcoxon Signed Ranks test was used for not normally distributed data. All statistical tests were performed on Graph Pad Prism.

