## Supplementary Figures for "Cycles of transcription and local translation support molecular long-term memory in the hippocampus"

### Supplementary Material:

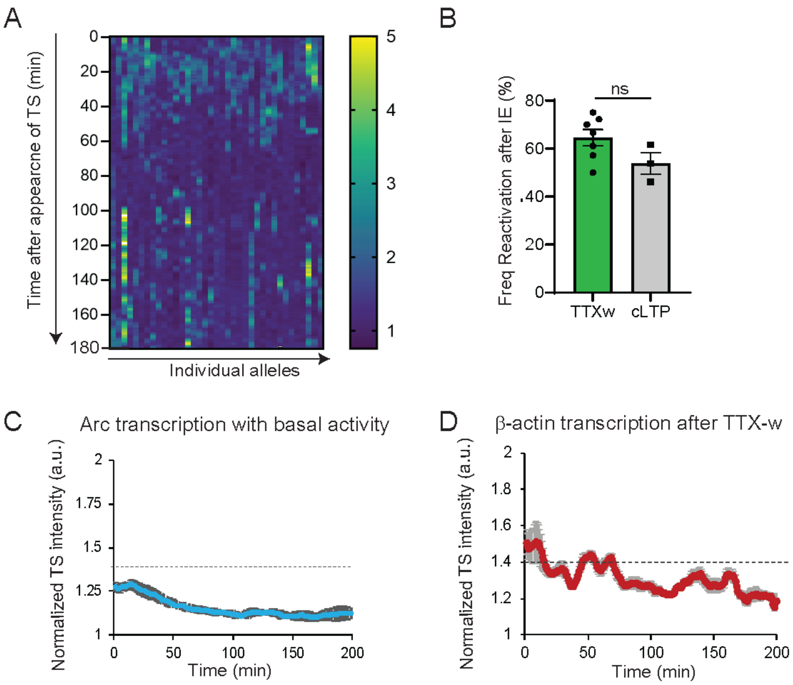


**Fig. S1.**

**Transcription profile of individual alleles reveals reactivation under different stimulation conditions.** (**A**) Heat map showing the transcription intensity of individual alleles over time after the appearance of transcription site (TS) post stimulation. Each column represents a single allele, and the rows represent time. The warmer colors indicate higher amplitude of transcription. Immediate early activation is followed by shutdown (darker colors) followed by reactivation (warmer colors). n = 37 alleles from 5 independent experiments. (**B**) Comparison of transcriptional reactivation frequency in activated neurons between two stimulation conditions. n = 92 cells from 6 experiments in TTX w, n = 34 cells from 3 independent experiments in cLTP. (**C**) Average intensity trace of Arc transcription from Arc-PBS animals monitored under basal network activity (no TTX added). n = 24 cells from 3 independent experiments. Biphasic transcription was not observed. (**D**) Average intensity trace of β-actin transcription from MBS-tagged β-actin allele after evoked activity upon TTX-withdrawal. Dashed line indicates threshold for inclusion criteria as active transcription. n = 20 cells from 3 independent experiments. Error bars indicate SEM.

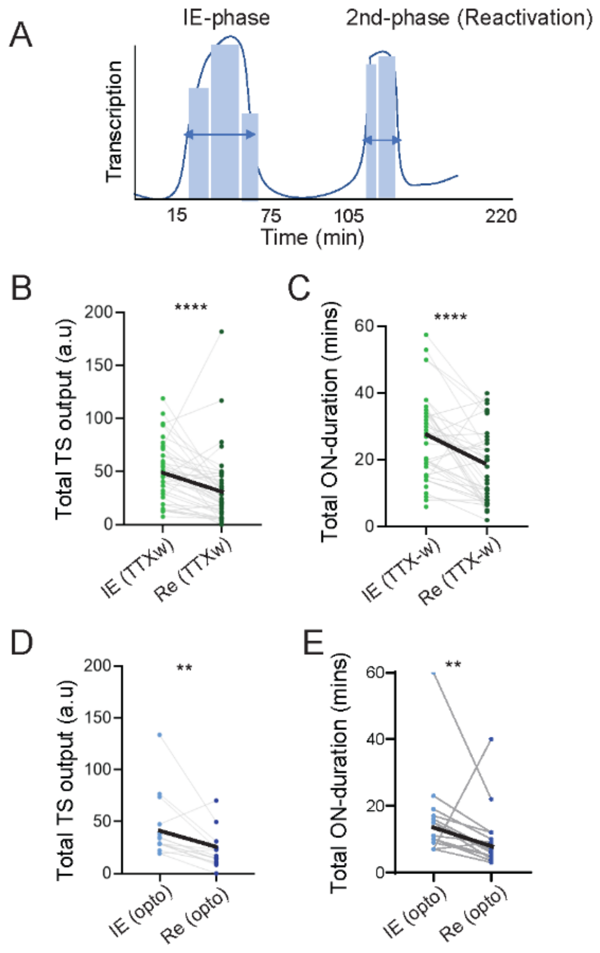


**Fig S2**

**Comparison of transcriptional output between immediate early and reactivation.** (**A**) Each transcriptional cycle can be composed of multiple bursts. The total ON-duration and the transcriptional output of all the bursts in the first cycle (IE; (15-75 min post stimulation), and the 2$^{nd}$ cycle (reactivation, Re; 105-200 min post stimulation) were calculated. (**B-E**) Pairwise comparison of total transcriptional output and ON-duration between the two cycles after TTX-w stimulation (B, C), and after single neuron stimulation (opto) (D,E). Each dot represents a single allele. n = 40 cells for TTX-w, n = 13 cells for opto, * $p < 0.05$, ** $p < 0.01$, **** $p < 0.001$.

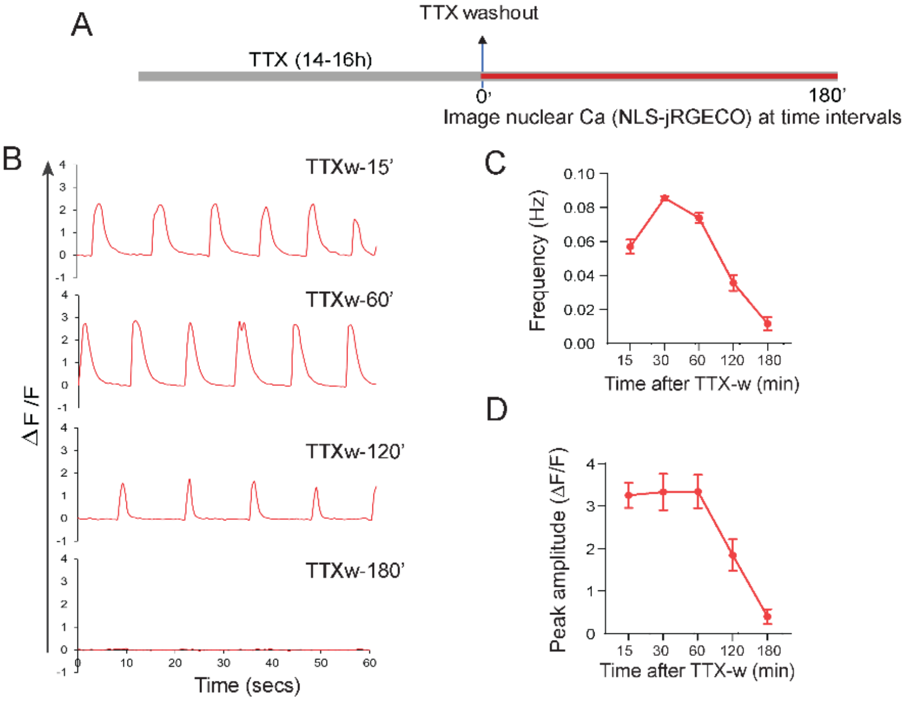


**Fig S3**:

**Nuclear calcium measurements over time.** (**A**) Schematic of stimulation paradigm and measurements of nuclear $Ca^{2+}$ levels using NLS-jRGECO1a. The same neurons were imaged at different time points after stimulation, where $Ca^{2+}$ activity was recorded for 1min at 1Hz acquisition rate. (**B**) Traces of nuclear $Ca^{2+}$ transients at different time points after TTXw. (**C**) Graphs showing frequency of $Ca^{2+}$ transients across time. (**D**) Graphs showing peak amplitude of $Ca^{2+}$ transients across time. n = 43 neurons from 3 independent experiments. Error bars indicate SEM.

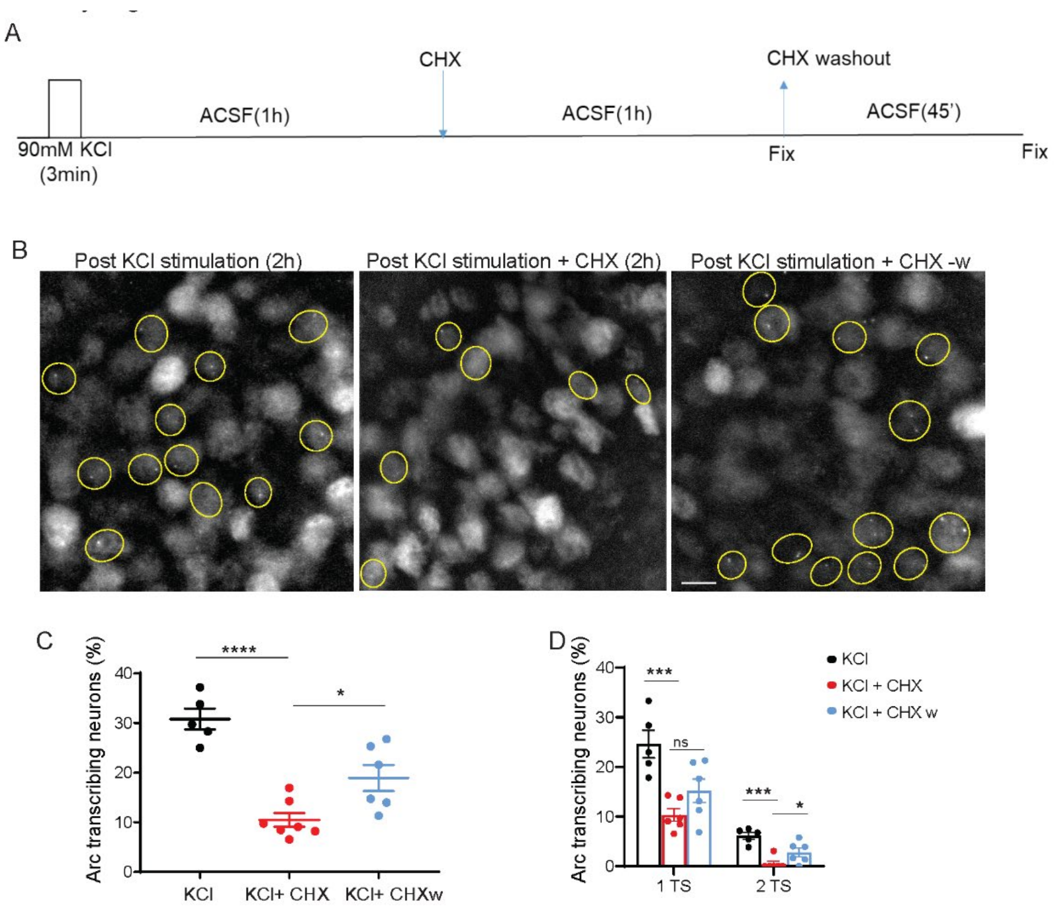


**Fig S4:**

**Protein synthesis dependent transcriptional phase in slices.** (**A**) Schematic of stimulation paradigm, where a brief (3 min) depolarization with KCl was performed followed by washout, and incubation of slices in ACSF. Acute hippocampal slices from Arc-PBS/PCP-GFP animals were used. Transcription was imaged *post hoc* after fixing the slices at different time points as shown. (**B**) Representative images of GC nuclei show distinct transcription sites at 2-hour post stimulation (*left panel*). Lack of transcribing cells with CHX incubation (*middle panel*) and restoration of transcription post CHX washout (*right panel*). Transcribing cells marked with yellow boundaries. (**C**) Percentage of transcribing neurons across different conditions (KCl vs KCl + CHX, p = , one-way ANOVA; KCl +CHX vs KCl + CHX-w, p= , one-way ANOVA . (**D**) Percentage of neurons exhibiting transcription from one or both alleles in different conditions. Note the absolute loss of neurons with 2 TS after CHX addition. (KCl vs KCl + CHX, p = , one-way ANOVA; KCl +CHX vs KCl + CHX-w, p= , one-way ANOVA. Error bars indicate SEM. n = 5 slices from 3 animals. * $p < 0.05$, ** $p < 0.01$, **** $p < 0.001$. Scale bar 10µm.

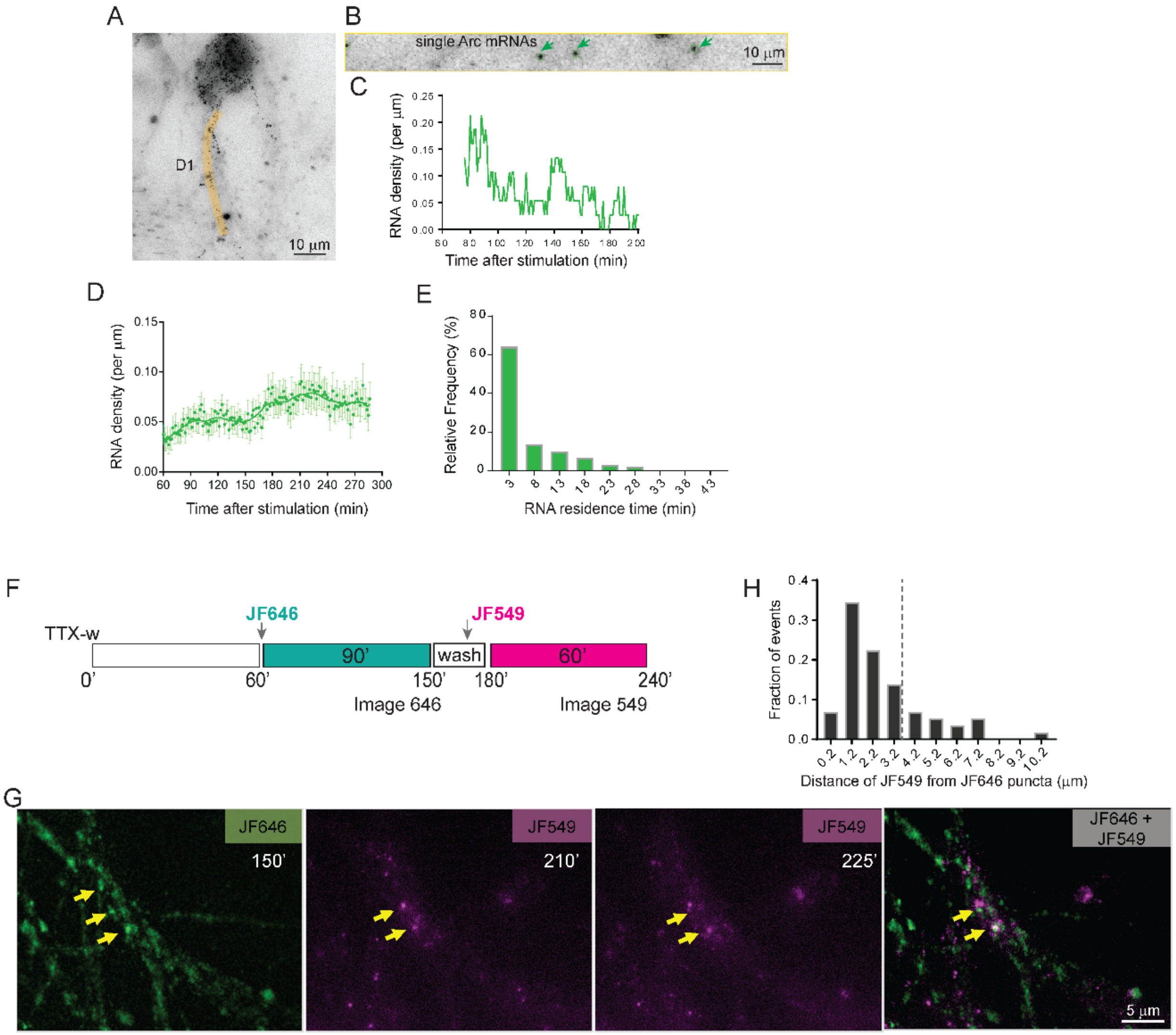


**Fig S5**:

**Long-term dynamics of *Arc* mRNAs and proteins in dendrites**. (**A**) Image of a neuron with Arc mRNAs in dendrites. Time lapse imaging of dendritic mRNAs performed for several hours after stimulation. (**B**) Image of a straightened dendrite (D1). Green arrows indicate single Arc mRNAs. (**C**) RNA density measured over time from the dendrite in (B). (**D**) Average mRNA density from multiple neurons shows two phases of RNA accumulation over time. Solid line indicates time averaging of 10 min. (**E**) Histogram of residence times of Arc mRNAs in the dendrites. (**F**) Labeling scheme to detect Arc proteins from early and late phases. (**G**) Representative images show JF646 and JF549 label in the same dendrite, and the overlay. Yellow arrows indicate co-localization or close proximity of JF646 and JF549 signal. (**H**) Histogram of distances between local maxima of JF646 puncta and the nearest JF549 puncta. Dashed line indicates the 75$^{th}$ percentile. Scale bar is 5 µm. n =14 dendrites for mRNAs from 3 independent experiments; n =10 neurons for proteins from 2 independent experiments.

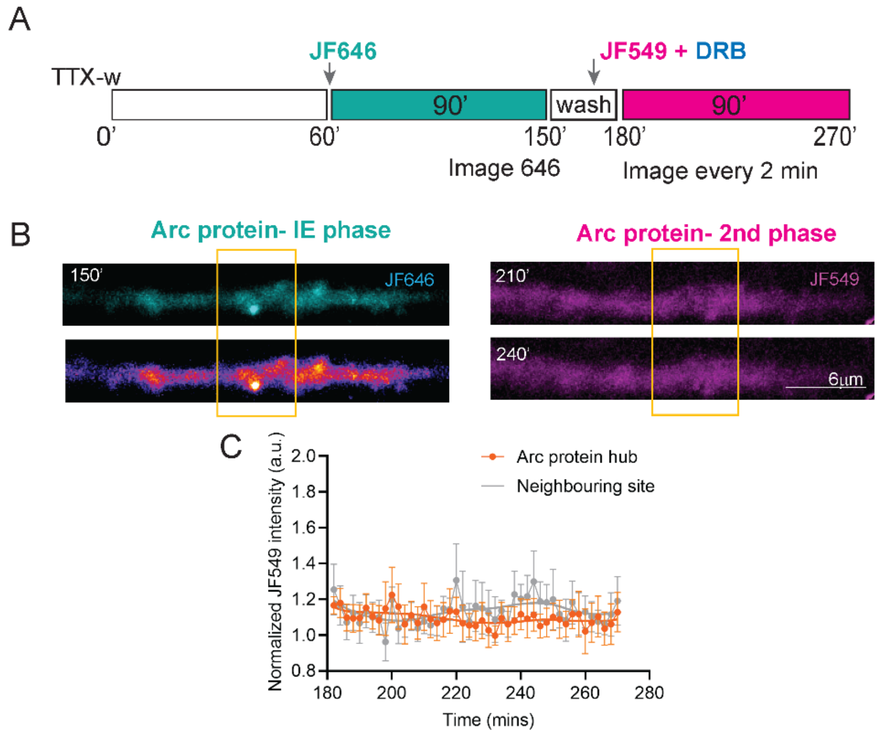


**Fig S6**:

**Inhibition of second transcription cycle disrupts Arc protein enrichment in hubs**. (**A**) Labeling scheme to detect Arc proteins from different phases, with addition of DRB after the IE-phase (165 min post stimulation). (**B**) Representative images show JF646 and JF549 label in the same dendrite. Lack of distinct JF549 puncta was observed with DRB treatment. (**C**) Normalized intensity trace of new Arc protein (JF549 signal) over time in the Arc hub versus in neighboring site as defined in Fig 5. Scale bar is 5 µm. n =10 dendrites from 2 independent experiments. Error bars show SEM.

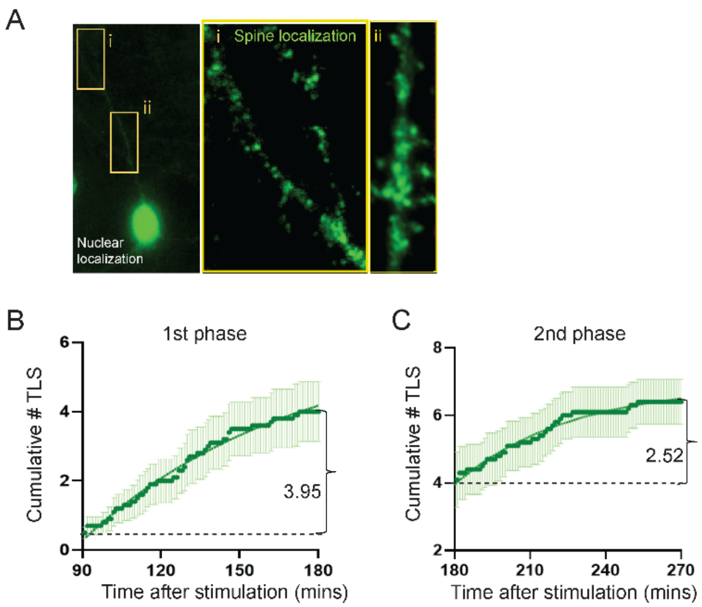


**Fig S7**:

**Arc translation reporter characterization and dynamics.** (**A**) Image of a neuron expressing the Suntag-Arc translation reporter, showing both nuclear and spine localization of the tagged Arc protein. Insets of dendrites have been adjusted for contrast to indicate spine localization of Suntag-Arc protein. (**B** Cumulative number of Arc TLS in dendritic hotspots from first phase (90-180 min), and second phase (180-270 min). Data fitted to a single-phase association model and change from baseline shown by brackets. n = 20 dendrites from 3 independent experiments.

**Supplementary Movies:**

**Movie S1**: Long term imaging of Arc transcription in neurons shows reactivation at the same allele at later time points (beyond IE-phase) after TTX-washout. Time in minutes.

**Movie S2**: Transcriptional cycling of the Arc gene at both alleles after TTX- withdrawal. Yellow and cyan arrows indicate the two different alleles in the same neuron. Time in minutes.

**Movie S3A**: Imaging of nuclear $Ca^{2+}$ transients in neurons using NLS-jRGECO1a after TTX-withdrawal and reapplication of TTX. Time in secs. The same neurons were followed over time, and $Ca^{2+}$ transients were completely abolished 20min post TTX application.

**Movie S3B**: Time lapse imaging of Arc transcription in neurons (from 3A) which were coexpressing PCP-GFP along with NLS-jRGECO1a after TTX-withdrawal and reapplication of TTX at 90 min. Yellow and cyan arrows indicate the two different alleles in the same neuron. Reactivation of Arc transcription observed even though CaTs were negligible (in Movie S3A). Time in mins.

**Movie S4**: Effect of translation inhibition on Arc transcription reactivation. Neurons expressing PCP-GFP were imaged over time after TTX-withdrawal, followed by addition of CHX, translation elongation inhibitor (50µg/ml) at 90 min, and then washout of CHX at 160min. Time in mins.

**Movie S5**: Time-lapse imaging of Arc mRNAs in dendrites for several hours after stimulation. Imaging was started at 75 min post TTX-withdrawal. Each green outline indicates a single Arc mRNA. Tracking show cycles of localization, with transient residence times.

**Movie S6**: Tracking of Arc translation sites in dendrites after stimulation. Primary hippocampal neurons were infected with Arc-Suntag reporter (in Fig 5A) and scFV-sfGFP for real time imaging of TLS (Fig 5B). The neuron was imaged everyone minute (time label in mins). Scale bar 5µm.
